# A novel drug-combination screen in zebrafish identifies epigenetic small molecule candidates for Duchenne muscular dystrophy

**DOI:** 10.1101/2020.02.19.956532

**Authors:** Gist H. Farr, Melanie Morris, Arianna Gomez, Thao Pham, Elizabeth U. Parker, Elisabeth Kilroy, Shery Said, Clarissa Henry, Lisa Maves

**Affiliations:** Center for Developmental Biology and Regenerative Medicine, Seattle Children’s Research Institute, Seattle, Washington, USA; Medical Student Research Training Program, University of Washington School of Medicine, Seattle, Washington, USA; Molecular Medicine and Mechanisms of Disease Program, Department of Pathology, University of Washington, Seattle, Washington, USA; Department of Pediatrics, University of Washington, Seattle, Washington, USA; School of Biology and Ecology, University of Maine, Orono, Maine, USA; University of Washington, Seattle, Washington, USA

## Abstract

Duchenne muscular dystrophy (DMD) is a severe neuromuscular disorder and is one of the most common muscular dystrophies. There are currently few effective therapies to treat the disease, although many small-molecule approaches are being pursued. Specific histone deacetylase inhibitors (HDACi) can ameliorate DMD phenotypes in mouse and zebrafish animal models and have also shown promise for DMD in clinical trials. However, beyond these HDACi, other classes of epigenetic small molecules have not been broadly and systematically studied for their benefits for DMD. Here, we performed a novel chemical screen of a library of epigenetic compounds using the zebrafish *dmd* model. We identified candidate pools of epigenetic compounds that improve skeletal muscle structure in *dmd* zebrafish. We then identified a specific combination of two drugs, oxamflatin and salermide, that significantly rescued *dmd* zebrafish skeletal muscle degeneration. Furthermore, we validated the effects of oxamflatin and salermide in an independent laboratory. Our results provide novel, effective methods for performing a combination small-molecule screen in zebrafish. Our results also add to the growing evidence that epigenetic small molecules may be promising candidates for treating DMD.

## Introduction

Duchenne muscular dystrophy (DMD) is a severe neuromuscular disorder caused by an X-linked mutation in the *DMD* gene, which encodes the protein dystrophin (Hoffman et al., 1987; Monaco et al., 1986). Dystrophin is a key component of the dystrophin-associated protein complex (DAPC), which serves as a stabilizing link between the cytoskeleton and extracellular matrix during muscle fiber contraction. Loss of dystrophin and DAPC function makes muscle cell membranes susceptible to contraction-induced damage and leads to progressive calcium dysregulation, satellite cell dysfunction, inflammation, fibrosis, and necrosis (reviewed in Allen et al., 2016 and in Spinazzola and Kunkel, 2016).

DMD is the most common type of muscular dystrophy, affecting approximately 1 in 3,500-5,000 male births (Emery, 1991; Mah et al., 2014). It first presents as motor difficulties in early childhood and progresses rapidly, leaving most affected boys in need of a wheelchair in their early teens and in need of respiratory aid in their 20s (reviewed in Crone and Mah, 2018). The disease is usually fatal in the third or fourth decade due to respiratory or cardiovascular failure.

Treatment options for DMD are still quite limited. The current standard of care is corticosteroid treatment, which delays the progression of muscle dysfunction but has serious side effects (Bushby et al., 2010; Crone and Mah, 2018; Kinnett et al., 2015). DMD gene therapy and gene editing approaches are very promising but face many challenges (Chamberlain and Chamberlain, 2017; Conboy et al., 2018; Duan, 2018; Min et al., 2019). Many small molecule approaches are being identified that could benefit DMD by modulating different pathological mechanisms downstream of the dystrophin mutation (Crone and Mah, 2018; Guiraud and Davies, 2017; Hoffman, 2019; Spinazzola and Kunkel, 2016). A current view in the DMD field is that a combination of therapies targeting different pathological mechanisms may ultimately be most beneficial for patients (Guiraud and Davies, 2017; Hoffman, 2019).

Histone deacetylase inhibitors (HDACi) are one class of small molecules that has shown promise for DMD in the *mdx* mouse, in *dmd* zebrafish, and in DMD clinical trials (Bettica et al., 2016; Johnson et al., 2013; Minetti et al., 2006; ClinicalTrials.gov NCT02851797, NCT03373968; reviewed in Consalvi et al., 2014). HDACi are examples of “epigenetic drugs”, small molecules that target chromatin modifications and transcriptional regulators. Histone acetylation, which is often linked with open chromatin and active transcription, is one example of an epigenetic modification and is the target of many HDACi (Kouzarides, 2007). There are four different classes of histone deacetylase (HDAC) proteins, with different functions and tissue expression patterns, and different HDACi often selectively inhibit specific HDACs (Xu et al., 2007). Studies in the *mdx* mouse have identified epigenetic components to the pathogenesis of DMD, including constitutive activation of HDAC2 and abnormal levels of certain histone modifications (Chang et al., 2018; Colussi et al., 2008; Colussi et al., 2009; Saccone et al., 2014). HDACi that have been shown to ameliorate pathology in the *mdx* mouse are the pan-HDACi Trichostatin A (TSA), valproic acid, phenylbutyrate, SAHA, and givinostat (ITF 2357), and the class I HDACi MS-275 (Colussi et al., 2010; Consalvi et al., 2011; Consalvi et al., 2013; Minetti et al., 2006; reviewed in Consalvi et al., 2014). TSA also ameliorates DMD in zebrafish (Bajanca and Vandel, 2017; Johnson et al., 2013), and givinostat is showing promise in DMD clinical trials (Bettica et al., 2016). However, beyond these HDACi, other classes of epigenetic small molecules have not been broadly and systematically studied for their benefits for DMD.

Zebrafish are an outstanding model for DMD, particularly for screening and evaluating novel drug therapies (Bassett et al., 2003; Bassett and Currie, 2004; Berger et al., 2010; Maves, 2014; Widrick et al., 2019). A zebrafish *dmd* mutant strain, also known as *sapje*, has a nonsense mutation in exon 4 and is a dystrophin-deficient model of DMD (Bassett et al., 2003; Berger et al., 2010; Granato et al., 1996). The zebrafish *dmd* mutation is autosomal recessive, and approximately 25% of the offspring from a heterozygous *dmd*+/- cross exhibit degenerative muscle lesions by about 3 days post-fertilization. *dmd* zebrafish exhibit many aspects of human DMD pathology; in particular, skeletal muscle fibrosis and inflammation, including infiltration of mononuclear cells (Bassett and Currie, 2004; Berger et al., 2010). *dmd* zebrafish offer several advantages for screening and evaluating drugs, including being amenable to rapid and high-throughput screening and affording an exceptional range of approaches for assessing treatment outcomes (Kawahara et al., 2011; reviewed in Maves, 2014 and in Widrick et al., 2019). Zebrafish eggs can be rapidly produced in large numbers, the resulting embryos readily absorb drug compounds, and muscle development and structure can easily be observed *in vivo* through birefringence techniques (Berger et al., 2012; Granato et al., 1996; Kawahara et al., 2011). Because the zebrafish *dmd* mutation, in contrast to the *mdx* mouse and DMD^*mdx*^ rat, is lethal during juvenile stages, survival can be used as an outcome measure for drug treatments (Hightower et al., 2020; Kawahara et al., 2011; Kawahara et al., 2014). Large-scale drug screens have highlighted the potential of *dmd* zebrafish for identifying new therapeutic compounds and targets as well as for understanding the molecular mechanisms behind DMD (Kawahara et al., 2011; Kawahara et al., 2014; Waugh et al., 2014). In addition, *dmd* zebrafish can replicate drug effects seen in *mdx* mice (Hightower et al., 2020; Johnson et al., 2013; Kawahara et al., 2011; Li et al., 2014; Winder et al., 2011). Because of this strong conservation, insight from zebrafish can inform mammalian DMD studies, while also taking advantage of the utility of high-throughput analysis.

Here we performed a pilot screen of the commercially-available Cayman Chemical Epigenetics Screening Library to identify additional small molecule epigenetic regulators that could improve the *dmd* zebrafish muscle phenotype. We identified candidate pools of epigenetic compounds that improve skeletal muscle structure in *dmd* zebrafish. We then identified a specific combination of two drugs, oxamflatin and salermide, that rescued *dmd* zebrafish skeletal muscle degeneration. Overall, our study provides novel and effective methods for performing a pooled drug-combination chemical screen and gives further evidence that epigenetic small molecules may be good candidates for successfully treating DMD.

## Materials and Methods

### Zebrafish husbandry

All experiments involving live zebrafish (*Danio rerio*) were carried out in compliance with the Institutional Animal Care and Use Committee guidelines of Seattle Children’s Research Institute and the University of Maine. Zebrafish were raised and staged as previously described (Westerfield, 2007). Time (hpf or dpf) refers to hours or days postfertilization at 28.5°C. The wild-type stock and genetic background used was AB. The zebrafish *dmd*^*ta222a*^ mutant line (also known as *sapje*) has been described and is a null allele (Bassett et al., 2003; Granato et al., 1996). *dmd*^*ta222a*^ genotyping was performed as previously described (Berger et al., 2011). Eggs were collected from about 20-30-minute spawning periods and raised in petri dishes in ICS water (300 mg Instant Ocean/L, 0.56 mM CaCl2, 1.2 mM NaHCO3) at 28.5°C.

### Small molecules

Epigenetic small molecule library screening was performed using the Cayman Chemical Epigenetics Screening Library (Item No. 11076, Batch No. 0455098). The Library was received as 10 mM stocks of each chemical in Dimethyl Sulfoxide (DMSO, Sigma). The composition of the Cayman Chemical Epigenetics Screening Library can vary. The version we obtained contained 94 chemicals distributed over two 96-well plates. The identities of the chemicals and plate lay-out are shown in Table 1. Additional chemicals were purchased individually from Cayman Chemical.

**Table 1.** Cayman Chemical Epigenetics Screening Library.

| Plate 1 |  | Plate 2 |  |
| --- | --- | --- | --- |
| Well | Contents | Well | Contents |
| A2 | 3-amino Benzamide | A2 | AGK2 |
| A3 | SB 939 | A3 | CAY10603 |
| A4 | PCI 34051 | A4 | Chaetocin |
| A5 | 4-iodo-SAHA | A5 | Splitomicin |
| A6 | Sirtinol | A6 | CBHA |
| A7 | C646 | A7 | M 344 |
| A8 | Ellagic Acid | A8 | Oxamflatin |
| A9 | Scriptaid | A9 | Salermide |
| A10 | Suberohydroxamic Acid | A10 | Mirin |
| A11 | Apicidin | A11 | Pimelic Diphenylamide 106 |
| B2 | UNC0321 (trifluoroacetate salt) | B2 | (S)-HDAC-42 |
| B3 | (-)-Neplanocin A | B3 | MS-275 |
| B4 | Cl-Amidine | B4 | HNHA |
| B5 | F-Amidine (trifluoroacetate salt) | B5 | RG-108 |
| B6 | JGB1741 | B6 | 2',3',5'-triacetyl-5-Azacytidine |
| B7 | UNC0638 | B7 | S-Adenosylhomocysteine |
| B8 | Phthalazinone pyrazole | B8 | UNC0224 |
| B9 | Isoliquiritigenin | B9 | Chidamide |
| B10 | CCG-100602 | B10 | 3-Deazaneplanocin A |
| B11 | CAY10669 | B11 | Sinefungin |
| C2 | Zebularine | C2 | Pyroxamide |
| C3 | Delphinidin chloride | C3 | WDR5-0103 |
| C4 | ITF 2357 | C4 | AMI-1 (sodium salt) |
| C5 | PFI-1 | C5 | UNC1215 |
| C6 | 5-Azacytidine | C6 | GSK 343 |
| C7 | Decitabine | C7 | SIRT1/2 Inhibitor IV |
| C8 | (+)-JQ1 | C8 | I-CBP112 (hydrochloride) |
| C9 | (-)-JQ1 | C9 | UNC1999 |
| C10 | BSI-201 | C10 | PFI-3 |
| C11 | 1-Naphthoic Acid | C11 | *trans*-Resveratrol |
| D2 | Sodium 4-Phenylbutyrate | D2 | 2,4-DPD |
| D3 | IOX1 | D3 | DMOG |
| D4 | MI-2 (hydrochloride) | D4 | Trichostatin A |
| D5 | MI-nc (hydrochloride) | D5 | CAY10398 |
| D6 | Gemcitabine | D6 | RSC-133 |
| D7 | Lomeguatrib | D7 | KD 5170 |
| D8 | Octyl-α-ketoglutarate | D8 | CAY10433 |
| D9 | Daminozide | D9 | Piceatannol |
| D10 | GSK-J1 (sodium salt) | D10 | CAY10591 |
| D11 | GSK-J2 (sodium salt) | D11 | EX-527 |
| E2 | GSK-J4 (hydrochloride) | E2 | SAHA |
| E3 | GSK-J5 (hydrochloride) | E3 | 2-PCPA (hydrochloride) |
| E4 | CPTH2 | E4 | Nicotinamide |
| E5 | Valproic Acid (sodium salt) | E5 | BIX01294 |
| E6 | Tenovin-1 | E6 | N-Oxalylglycine |
| E7 | Tenovin-6 | E7 | Suramin (sodium salt) |
| E8 | Sodium Butyrate |  |  |
| E9 | Anacardic Acid |  |  |

### Drug treatments

To test toxicity and assess working doses of the Epigenetics Screening Library chemicals, wild-type AB embryos were treated with individual drugs beginning at 4 hpf. Embryos were not dechorionated prior to these dose tests. Three concentrations of each drug were tested: 10 µM, 1 µM, and 100 nM. Embryos were treated with drugs with 1% DMSO in embryo medium (EM), or with 1% DMSO in EM as a vehicle control. 12-well plates were used; each well contained 25 embryos and 3 mL of drug treatment. The drug treatment was changed every 24 hours until 4 dpf. Over the 4-day treatments, developmental abnormalities and survival were noted. Most drugs showed little or no effect on embryo health and survival at 1 µM doses, so we determined 1 µM to be the working dose for drug pool screening for most of the Library drugs. C646 and Apicidin were toxic at 1 µM but not at 100 nM. CAY10669, GSK-J1, GSK-J4, and Tenovin-1 exhibited strong toxicity and thus were removed from drug pool screening.

For drug pool screening and *dmd* rescue tests, embryos from *dmd+/-* crosses were used. At about 24 hpf, the chorions were manually removed from the embryos, and embryos were sorted into fresh petri dishes or wells for drug treatments. 1000X stocks of drugs were made in DMSO just prior to each drug treatment experiment. Embryos were treated with drugs with 1% DMSO in embryo medium (EM), or with 1% DMSO in EM as a vehicle control. For the drug pool screen, 25 embryos each were sorted into two wells of a 12-well plate and treated with 3 mL of EM containing either the drug treatments or vehicle control. Treatments were started at about 24 hpf and continued for 3 days, changing the drug treatment or vehicle control EM solution every 24 hours. Larvae were fixed in 4% paraformaldehyde (PFA) at 4 dpf and stored at 4°C. Larval heads were removed for genotyping, and tails were maintained in 4% PFA.

### Imaging and scoring muscle lesions

Larvae fixed in 4% PFA were rinsed in PBS with 0.025% Tween (PBSTw) prior to imaging. Animals were placed in PBSTw in a 60 mm glass Petri dish. An Olympus SZX16 stereomicroscope with attached Olympus DP72 camera was set up with one sheet of polarizing film over the trans-illumination base and another sheet of the film placed over the objective lens such that the two films were crossed. For drug pool screening, larvae were sorted under birefringence into muscle lesion positive or negative groups. The number of larvae in each group was recorded.

For quantitative birefringence measurements, we developed an approach inspired by a previously described method (Berger et al., 2012). In our approach, larvae were placed in a glass-bottom petri dish in 2.5% methyl cellulose and oriented to maximize the brightness of the muscle tissue through the crossed polarizers. Birefringence was imaged in individual animals as above. Images were acquired using Olympus Cellsens Dimensions software. Exposure time was adjusted so that only a few saturated pixels were present. ImageJ was used to outline the trunk musculature, using the wand tool, and to calculate the average pixel intensity within the resulting selection.

### Statistical analyses

Statistical tests used are provided in the Figure Legends. Statistical analyses were performed and graphs were constructed using GraphPad Prism 8.

## Results

### Epigenetic small-molecule library pooling approach

In a previous study, we showed that the pan-HDACi TSA could improve the zebrafish *dmd* muscle lesion phenotype (Johnson et al, 2013). We wanted to test whether we could identify new epigenetic drugs, or epigenetic drug combinations, that improve the zebrafish *dmd* muscle lesion phenotype. We turned to the Cayman Chemical Epigenetics Screening Library. The version of the Library that we obtained contained 94 chemicals distributed over two 96-well plates (Table 1). In order to efficiently test combinations of drugs in the Library, we designed a grid system consisting of a set of 393 different drug pools (Supplemental Table 1). In this system, drug A2 of Plate 1 is pooled with the remaining drugs of Row A of Plate 1 for one combination pool. Then drug A2 is pooled with Row B drugs of Plate 1 for a second combination pool, and then drug A2 is pooled with the remaining rows of drugs on both Plate 1 and Plate 2. Drug A3 of Plate 1 is then combined with drugs of Row B of Plate 1, and so on (Supplemental Table 1). In this grid system, each drug would be tested in combination with each of the other Library drugs in at least 1 pool (Supplemental Table 1).

Prior to screening these chemicals on *dmd* embryos, we performed dose and toxicity testing of each individual Library drug on wild-type embryos (see Materials and Methods). Most drugs showed little or no detrimental effect on embryo health and survival at 1 µM dose, so we selected 1 µM to be the working dose for drug screening for most of the Library drugs. Four drugs exhibited strong toxicity, so we removed these from the drug pool grid (CAY10669, GSK-J1, GSK-J4, and Tenovin-1; Table 1; Supplemental Table 1). We also removed Library drugs that are described as negative control drugs from the drug pool grid ((-)-JQ1, MI-nc, GSK-J2, and GSK-J5; Table 1; Supplemental Table 1).

To determine if screening pools of epigenetic small molecules was sensitive enough to identify drugs that could improve the zebrafish *dmd* muscle phenotype, we tested the pool of 10 drugs from Plate 2 Row D, which contains TSA (Table 1). Animals from *dmd+/-* crosses were treated from 1-4 days and then scored for muscle birefringence (as in Johnson et al., 2013 and illustrated in Figure 1). Treatment with the pool of drugs from Plate 2 Row D significantly decreased the number of affected animals exhibiting abnormal muscle birefringence (Figure 2). These results indicate that a pool of drugs containing a known beneficial epigenetic drug, TSA, can improve the zebrafish *dmd* muscle phenotype. These results suggest that the epigenetic drug-pooling design of our screen is sensitive enough to pick up positive hits of small molecules that improve the zebrafish *dmd* muscle phenotype.

**Figure 1.**
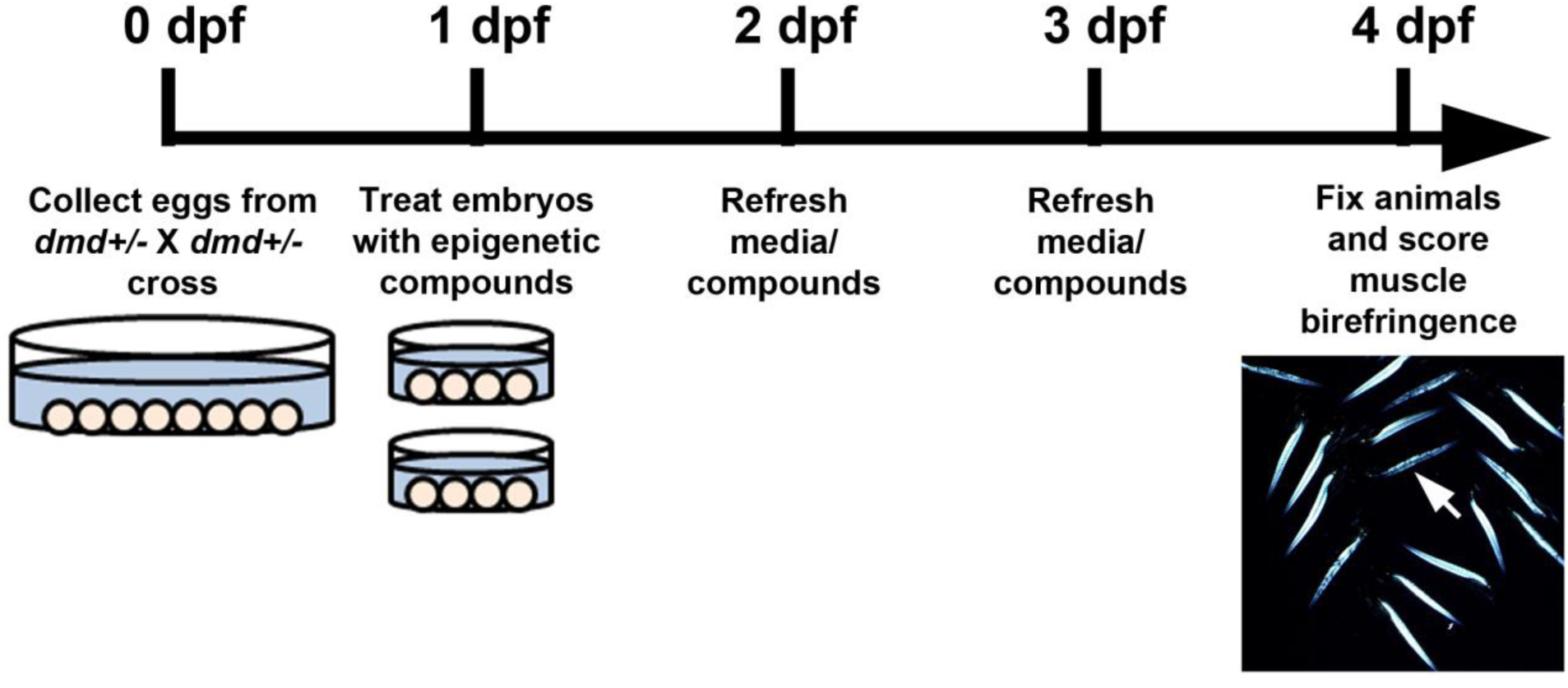
Timeline of *dmd* zebrafish drug treatments. Eggs are collected from crosses of *dmd+/-* fish. *dmd* is not sex-linked in zebrafish, so about 25% of embryos will be *dmd-/-*. At 1 dpf (days post fertilization), embryos are sorted into dishes for treatments, and DMSO or small molecule compounds are added to the embryo bath (as in Johnson et al., 2013). Drugs are replaced daily. At 4 dpf, animals are fixed and muscle birefringence is scored. *dmd-/-* animals exhibit dark lesions in the larval trunk muscle, as visualized using polarized light birefringence (arrow in image).

**Figure 2.**
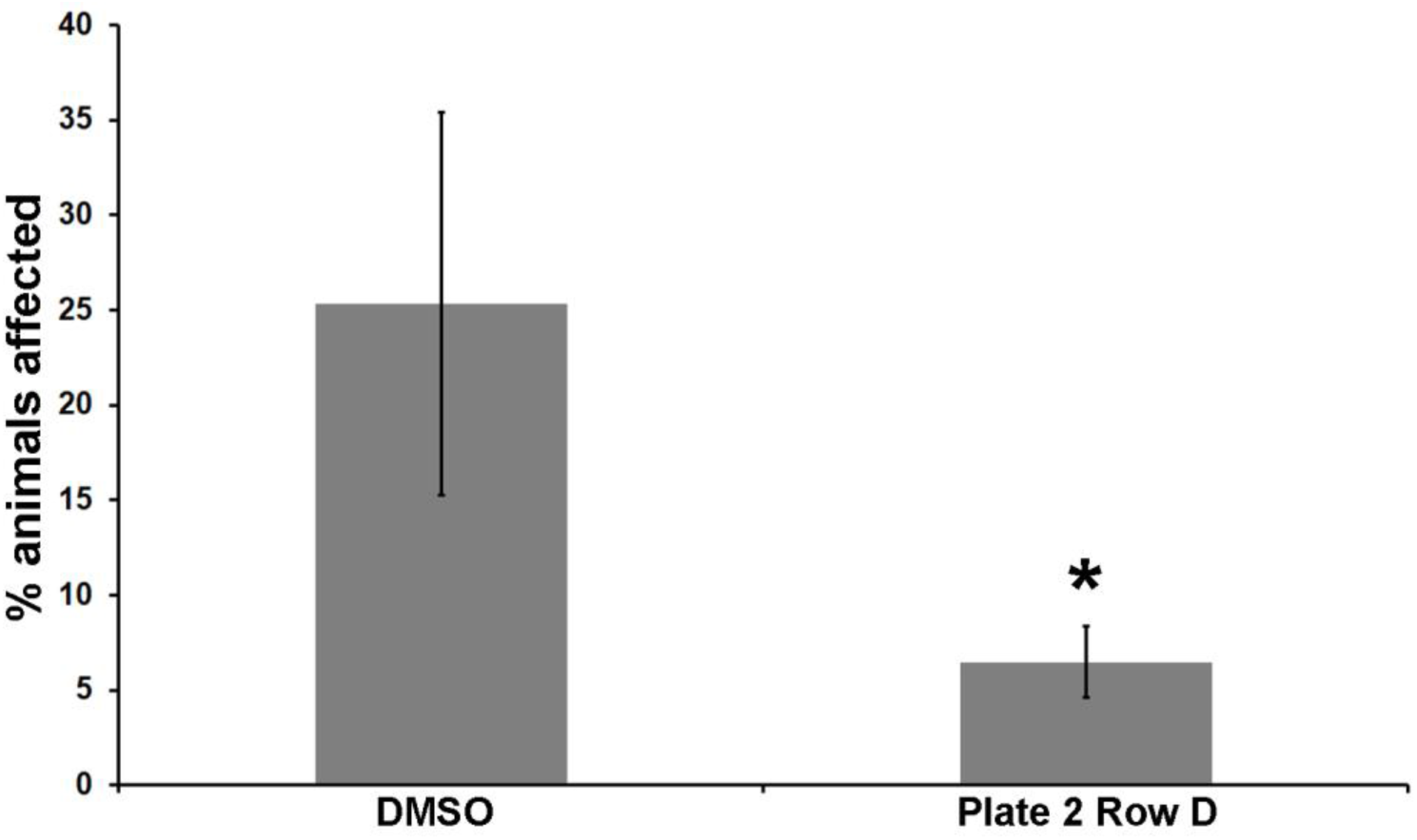
Plate 2 Row D drug pool, which includes TSA, significantly reduces the frequency of *dmd* zebrafish with abnormal muscle birefringence. Control treatment is 1% DMSO. See Table 1 for the 10 drugs in Plate 2 Row D. Each drug was used at 1 µM. For each treatment condition, n=3 replicates, with 23-27 embryos in each replicate. Error bars represent standard deviation. Significance was determined using a Mantel-Haenszel test. *, *p* < 0.004 compared to DMSO control.

### Pilot screen of epigenetic small-molecule combination pools

We then proceeded to test epigenetic drug pools from our drug-combination grid system (Supplemental Table 1). We tested 93 of the 393 possible drug pool combinations (Figure 3A; Supplemental Table 1). As above, animals from *dmd+/-* crosses were treated from 1-4 days and then scored for muscle birefringence. Each pool was tested in duplicate. We determined the average percentage of affected animals from each drug pool, and from each DMSO control tested, and we plotted the results as a heat map on our drug pool grid (Figure 3A). This heat map grid reveals that, while no specific individual drug well appears to have a strong effect on reducing the percentage of animals affected (horizontal rows in Figure 3A), the majority of pools tested that included Plate 2 Row A reduce the percentage of animals affected (vertical column 5 in Figure 3A). To test whether the Plate 2 Row A pools are significantly improving the percentage of animals affected, we plotted the combined results from all pools containing each plate row compared with the combined results from DMSO control treatments (Figure 3B). This analysis shows that treatment pools using Plate 2 Row A led to a significant reduction in the percentage of affected animals relative to the DMSO controls (Figure 3B). On average, treatment with drug pools that included Plate 2 Row A resulted in only about 10% affected animals, compared to the 25% affected average observed in the DMSO controls (Figure 3B). Figure 3C-D shows an example of the improved birefringence images from animals treated with a representative drug pool from the Plate 2 Row A pools (Pool 59), compared to sibling DMSO controls. While these results do not rule out beneficial effects of compounds or treatment pools from our screen that did not include Plate 2 Row A, these results do highlight that compounds within Plate 2 Row A are having significant beneficial effects on *dmd* zebrafish.

**Figure 3.**
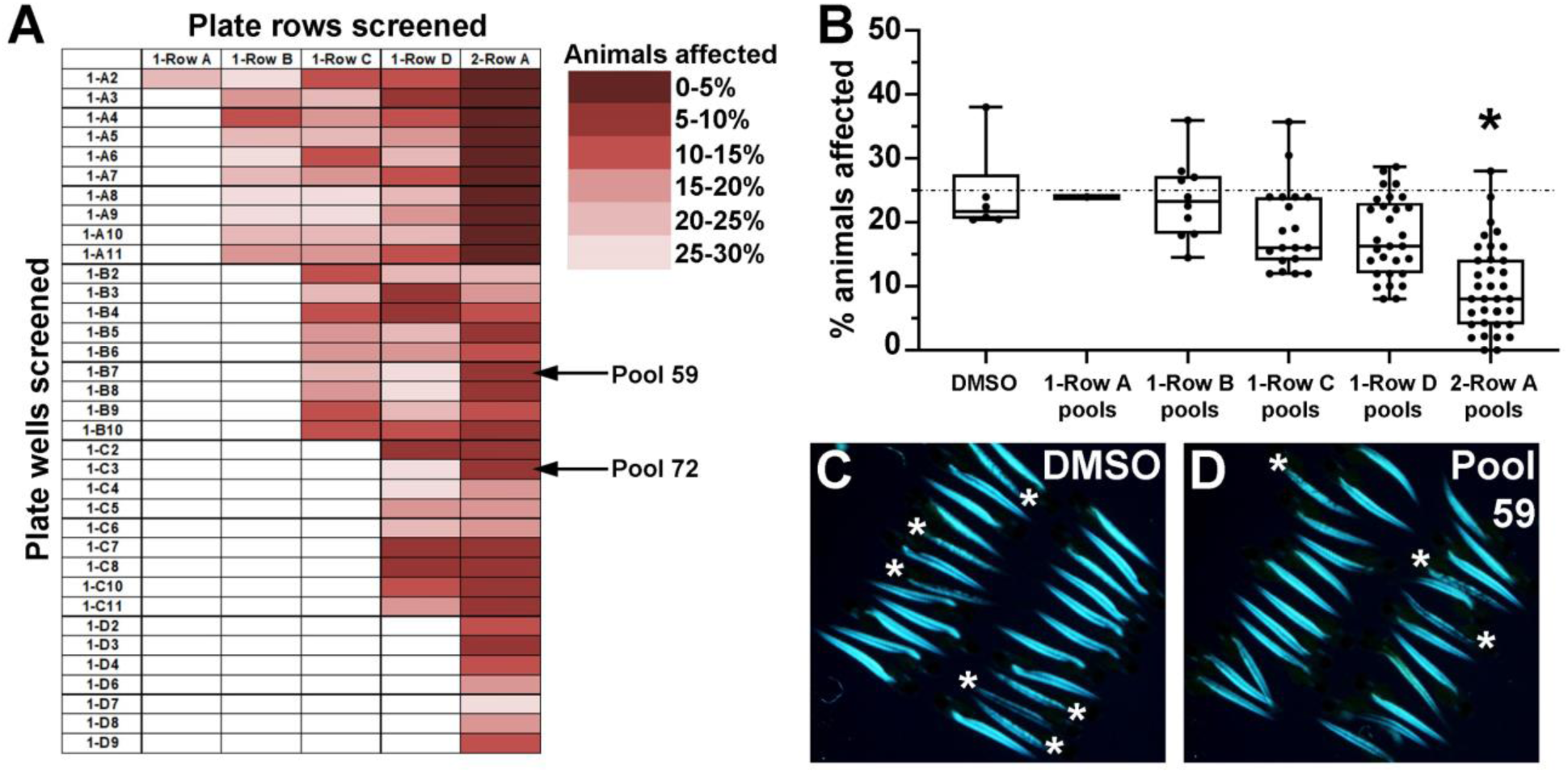
Pilot screen highlights the ability of Plate 2 Row A drug pools to lower the frequency of affected *dmd* animals. (A) Heat map representation of the average percentage of affected animals observed from each of the 93 drug pools tested. Individual drug wells tested were from Plate 1 (left column, plate number and well number shown). Drug library rows tested are Rows A-D from Plate 1 and Row A from Plate 2. See Table 1 for the drugs in each well and row. Arrows point to Pool 59, shown in (D), and Pool 72, analyzed in Figure 4 below. (B) Graph of combined average percentages of affected animals from all tested pools that included each plate row. Control treatments are 1% DMSO. The dashed line represents the average of the control DMSO treatments (25%). Boxes extend from the 25^th^ to the 75^th^ percentile, the median is denoted with a bar, and the whiskers extend to the highest and lowest values. Significance was determined using a one-way ANOVA test comparing each drug pool group to the DMSO control group. Dunnett’s test was used to correct for multiple comparisons. *, *p*=0.0021 compared to DMSO control. (C-D) Example of birefringence images of 4 dpf larvae from a single replicate of (C) DMSO control and (D) Pool 59 (Plate 2 Row A + UNC0638; D) treatments. In this example, DMSO animals are siblings of Pool 59 animals. Asterisks mark affected animals.

### Analysis of Plate 2 Row A drugs identifies the beneficial combination of oxamflatin and salermide

In order to determine which drug or drugs are mediating the effect of Plate 2 Row A, we wanted an approach that would allow us to more quantitatively assess the effects of drugs on *dmd* birefringence. We developed a method that uses gray value measurements to quantitate the brightness level of the muscle birefringence of each animal (Figure 4A). For each animal, we outlined a unilateral area of trunk muscle birefringence and measured the average pixel brightness within that area (Figure 4A). *dmd* animals show lower pixel intensity than their control siblings (Figure 4B,E). As a test of this approach, we confirmed that TSA treatment significantly improved the average pixel brightness of *dmd* birefringence (Figure 4D-E).

**Figure 4.**
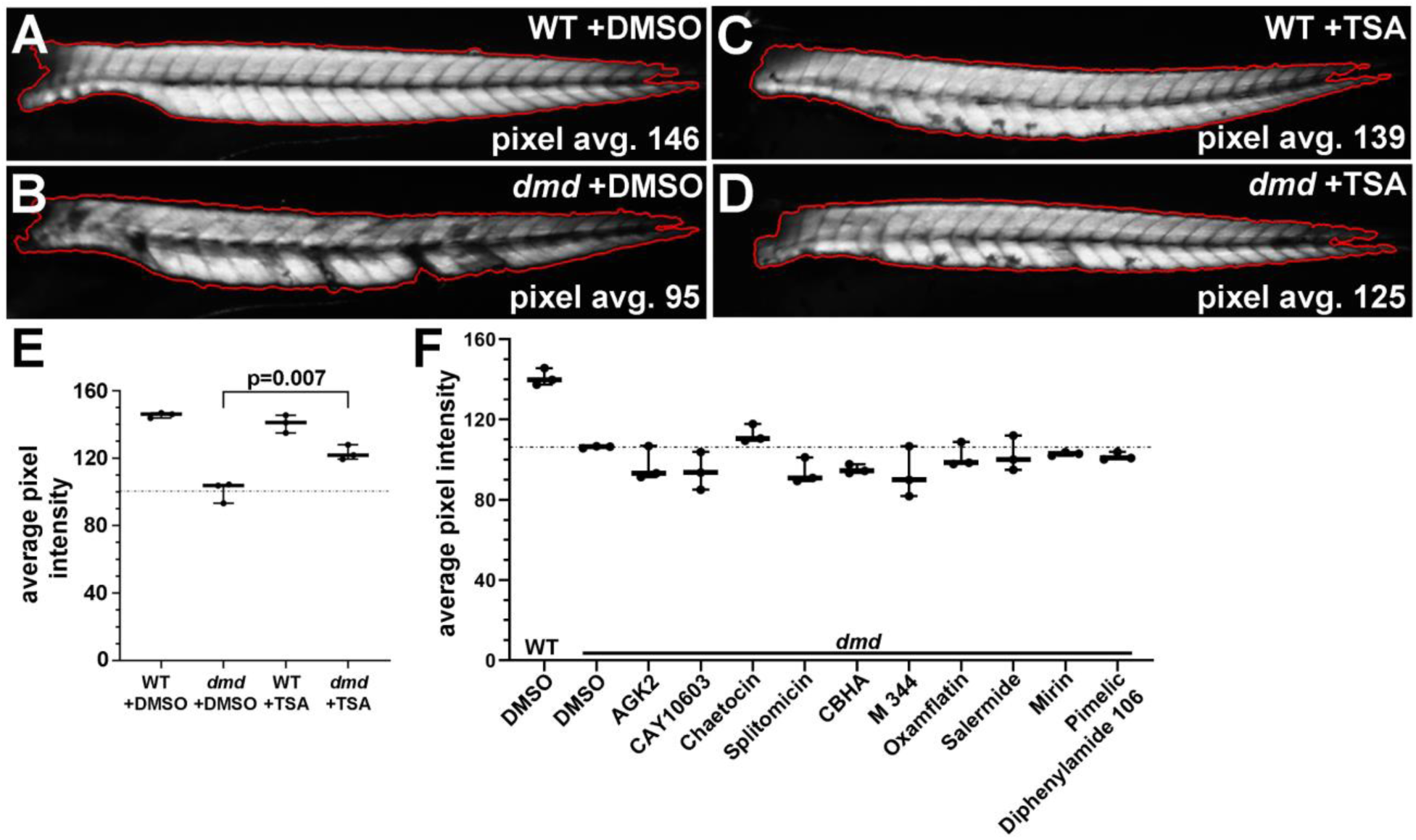
Individual drugs from Plate 2 Row A do not significantly improve *dmd* zebrafish muscle birefringence. (A-D) 4-day-old zebrafish trunk muscle, with trunk skeletal muscle birefringence outlined in red. Anterior to the left. Average pixel brightness values are shown for wild-type (WT) +DMSO (A), *dmd* +DMSO (B), WT +TSA (C), and *dmd* +TSA (D). (E) Graph of average pixel intensities for 400 nM TSA treatments vs DMSO controls. n=3 biological replicates (25-embryo pools) for each treatment. Plot shows the average pixel intensity for each of the 3 replicate pools for each treatment (3-13 genotyped animals per pool). The dashed line represents the average pixel intensity for all of the DMSO-treated *dmd* animals (n=14). P value determined by Student’s t-test. (F) Graph of average pixel intensities for treatments of *dmd* fish with the 10 individual drugs from Plate 2 Row A. Control treatment is 1% DMSO. All drugs were tested at 1 µM. For each treatment condition, n=3 replicates, with 2-9 *dmd* embryos in each replicate. Plot shows the average pixel intensity for each of the 3 replicate pools for each treatment. The dashed line represents the average pixel intensity for all of the vehicle-treated *dmd* animals (n=15). Significance was determined using a one-way ANOVA test comparing each treatment group to the *dmd* DMSO control group. Dunnett’s test was used to correct for multiple comparisons. *p*>0.2 for all individual drugs compared to *dmd* DMSO control.

We next tested each drug from Plate 2 Row A to see if any of the individual drugs could improve the *dmd* birefringence phenotype. Animals from *dmd+/-* crosses were treated from 1-4 days and then scored for muscle birefringence. When tested individually, none of the drugs showed a significant effect on the DMSO-treated *dmd* muscle birefringence brightness (Figure 4F). These results indicate that a combination of drugs from Plate 2 Row A is likely needed to improve the *dmd* birefringence phenotype.

We then tested for drugs required for the effects of Plate 2 Row A by removing individual drugs. We decided to test drug screen Pool 72 (Plate 2 Row A + Plate 1 Well C3; 11 total drugs), because this pool represented an average effect from the Plate 2 Row A pools (Figure 3A-3B). As above, animals from *dmd+/-* crosses were treated from 1-4 days and then scored for muscle birefringence. Treatment with the full 11 drugs from Pool 72 showed significant improvement of the *dmd* birefringence using our quantitative analysis (Figure 5A). However, the removal of four different drugs (chaetocin, oxamflatin, salermide, and delphinidin chloride) each inhibited the ability of Pool 72 to improve *dmd* birefringence (Figure 5A), indicating that these four drugs may be involved in producing the *dmd* rescue effect of Pool 72.

**Figure 5.**
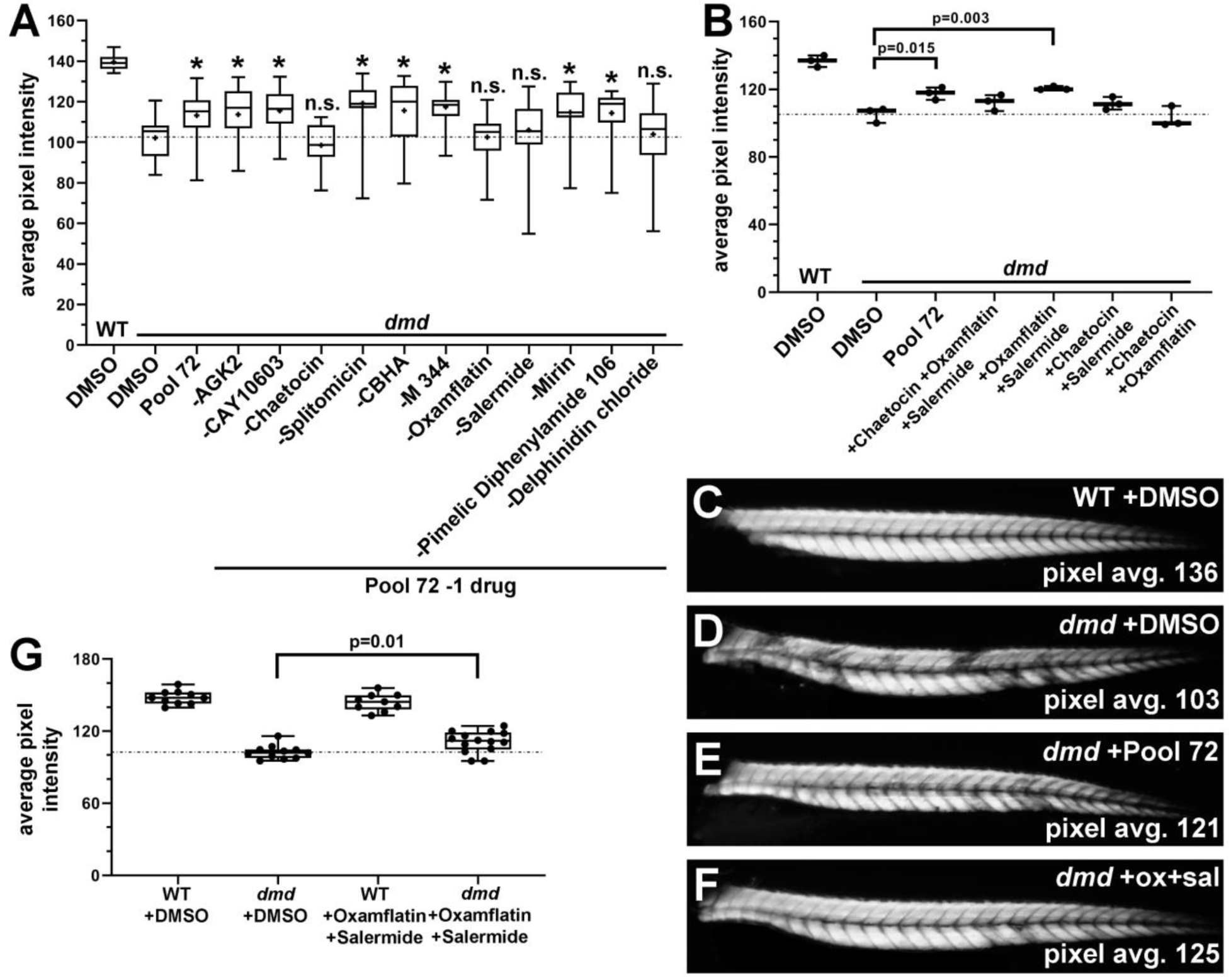
Oxamflatin and salermide mediate the effects of Plate 2 Row A. (A) Graph of average pixel intensities for treatments of *dmd* fish with Pool 72 and with Pool 72 with each individual drug removed. Control treatment is 1% DMSO. All drugs included in the pools were tested at 1 µM. For each condition, 3 pools of 25 embryos were treated, with 2-9 *dmd* embryos in each pool. The boxes on the plot represent the average pixel intensities of the individual *dmd* embryos from all 3 pools for that treatment (n=14-26 total *dmd* embryos per treatment). Treatment with Pool 72 without chaetocin, oxamflatin, salermide, or delphinidin chloride did not achieve the rescue effect seen with the full Pool 72. Boxes extend from the 25^th^ to the 75^th^ percentile, the median is denoted with a bar, the mean is indicated with a cross, and the whiskers extend to the highest and lowest values. The dashed line represents the average pixel intensity for all of the DMSO-treated *dmd* animals (n=26). Significance was determined using a one-way ANOVA test comparing each treatment group to the *dmd* DMSO control group. Dunnett’s test was used to correct for multiple comparisons. *, *p*≤0.04 compared to *dmd* DMSO control. (B) Graph of average pixel intensities for treatments of *dmd* fish with Pool 72 and with combinations of chaetocin, oxamflatin, and salermide. Control treatment is 1% DMSO. All drugs were used at 1 µM. For each treatment condition, n=3 replicates, with 1-11 *dmd* embryos in each replicate. Plot shows the average pixel intensity for each of the 3 replicate pools for each treatment. The dashed line represents the average pixel intensity for all of the DMSO-treated *dmd* animals (n=16). Significance was determined using a one-way ANOVA test comparing each treatment group to the *dmd* DMSO control group. Dunnett’s test was used to correct for multiple comparisons. (C-F) 4-day-old zebrafish trunk muscle birefringence. Anterior to the left. Representative animals are shown from drug treatments in (B). Average pixel brightness values are shown for (C) control, (D) *dmd* +DMSO, (E) *dmd* +Pool 72, and (F) *dmd* +oxamflatin and salermide. (G) Validation of oxamflatin and salermide treatment effects. Graph of average pixel intensities for treatments of *dmd* fish with oxamflatin and salermide, performed in the Henry Lab. Treatments were performed from 1 dpf-4 dpf. Control treatment is 1% DMSO. Drugs were each used at 1 µM. All animals were genotyped. Filled circles on the graph represent each individual animal (n=9-14). Boxes extend from the 25^th^ to the 75^th^ percentile, the median is denoted with a bar, and the whiskers extend to the highest and lowest values. Dashed line represents the average pixel intensity for all of the DMSO-treated *dmd* animals (n=11). P value determined by Student’s t-test.

We next focused on the roles of chaetocin, oxamflatin, and salermide. We decided to not further test the role of delphinidin chloride because it was not a component of the Plate 2 Row A drugs and because it did not exhibit beneficial activity on its own (data not shown). To further test the roles of chaetocin, oxamflatin, and salermide in improving *dmd* muscle birefringence, we tested these three drugs together and each pair-wise combination (Figure 5B). The combination of oxamflatin and salermide significantly improved *dmd* muscle birefringence, similar to Pool 72, whereas the other pair-wise combinations and the three-drug combination did not (Figure 5B-F). These results show that the combination of oxamflatin and salermide mediate the beneficial effects of Pool 72 on improving *dmd* muscle birefringence.

### An independent laboratory test validates the beneficial effects of oxamflatin+salermide

To validate these effects of oxamflatin and salermide, we performed treatments from 1 dpf-4 dpf at an independent site, in the laboratory of Dr. Henry. We see similar improvement of *dmd* muscle birefringence with oxamflatin+salermide treatments performed in the Henry Lab (Figure 5G) as we do with treatments performed in the Maves Lab (Figure 5B). This independent test is a strong validation of the benefits of this drug combination for *dmd* zebrafish.

## Discussion

In this study, we performed a screen of epigenetic-modifying small molecules to identify compounds that prevented muscle degeneration in the zebrafish model of DMD. We developed a novel drug-pooling approach to screen combinations of epigenetic drugs, and we performed a pilot screen of 93 novel epigenetic drug pools. While our pilot screen identified several new candidate drug pools that have beneficial effects on *dmd* zebrafish (Figure 3A), our drug-pooling strategy and subsequent analysis highlighted the activity of drugs from Plate 2 Row A (Figure 3). We were able to resolve a novel drug combination, oxamflatin and salermide, that significantly reduced the number of *dmd* animals with muscle lesions when compared to DMSO-treated controls. Our results provide support for continued efforts in screening epigenetic small molecules for their beneficial effects in *dmd* zebrafish, and they support further investigations of oxamflatin and salermide as potential therapeutic compounds for DMD.

The development of significantly beneficial therapies for DMD will likely involve the simultaneous use of a combination of drugs that target dystrophin and/or downstream pathological mechanisms (Guiraud and Davies, 2017; Hoffman, 2019). Epigenetic drugs may represent a promising component of a DMD combination therapy. Epigenetic drugs are outstanding candidates for a DMD pharmacological therapy for many reasons. Certain epigenetic drugs have been shown to benefit DMD and other neuromuscular disorders (Campbell et al., 2017; Consalvi et al., 2014; Gordon et al., 2013; Riessland et al., 2010). Several HDACi, including SAHA, and other epigenetic drugs are already FDA-approved for use in cancers (Abdelfatah et al., 2016; Ahuja et al., 2016; Bai et al., 2019). While some epigenetic drugs have off-target or solubility issues, there is potential for optimizing dosing or using these drugs as lead compounds for further analyses (Abdelfatah et al., 2016; Bai et al., 2019). In addition, there are growing numbers of examples showing that epigenetic small molecules synergize with each other or with other drugs to target different diseases, including cancer and heart disease (Abdelfatah et al., 2016; Ahuja et al., 2016; Alexanian et al., 2019; Bai et al., 2019; Huber et al., 2011; Mazur et al., 2015). In our work here, we show an example of a combination of two epigenetic drugs that are beneficial for *dmd* zebrafish. Because of the advantages of the zebrafish animal model, we expect that *dmd* zebrafish will be an increasingly important model for investigating combination therapies for DMD.

Oxamflatin is an HDACi that inhibits Class I and II HDACs and is chemically similar to TSA (Huber et al., 2011; Kim et al., 1999). Salermide is a Class III HDACi that inhibits the NAD+-dependent deacetylases SIRT1 and SIRT2 (Lara et al., 2009), and so represents a new class of HDACi for DMD. In previous studies, overexpression of SIRT1 has been shown to ameliorate DMD pathophysiology in *mdx* mice (Chalkiadaki et al., 2014), while salermide inhibition of SIRT1 has been shown to protect muscle cells against oculopharyngeal muscular dystrophy in *Caenorhabditis elegans* (Pasco et al., 2010). SIRT1 and SIRT2 can have opposing effects in promoting angiogenesis and in providing neuroprotective effects in neurodegenerative disease models (Keskin-Aktan et al., 2018; Shi et al., 2014). It will be important to determine the mechanisms by which salermide and oxamflatin are together providing skeletal muscle benefits in *dmd* zebrafish.

In conclusion, we developed a novel approach for screening combinations of small molecules in *dmd* zebrafish, and we identified a promising new combination of epigenetic compounds, oxamflatin and salermide. Future studies will include screening larger libraries of epigenetic small molecules in *dmd* zebrafish and pursuing validations of additional candidate drug pools that were identified in our pilot screen. Also, a critical next step will be to validate the effects of oxamflatin and salermide in mammalian DMD models, such as *mdx* mice, the *Dmd* rat, or human induced pluripotent stem cell models of DMD (Choi et al., 2016; reviewed in McGreevy et al., 2015; and Wells, 2018). In this work, we have already taken an important step in small molecule validation for DMD by demonstrating that oxamflatin and salermide show beneficial effects on *dmd* zebrafish in two independent laboratories. We predict that small molecule validation in independent laboratories and in multiple model systems will increase the potential for future translation of epigenetic drug therapies and other pharmacological approaches for DMD.

## Acknowledgements

We would like to thank the SCRI Office of Animal Care for caring for the zebrafish, and Kimia Imani, Max Urbanek, and Ermias Yohannes for their assistance. This work was supported by grants 1R03AR065760 (NIH, NIAMS), 1R21AR068536 (NIH, NIAMS), and MDBR-15-107-DMD (Million Dollar Bike Ride Grant Program) to L.M. Finally, we would like to thank the University of Washington School of Medicine Medical Student Research Training Program for financial support.

## Figures

**Supplemental Table 1.**
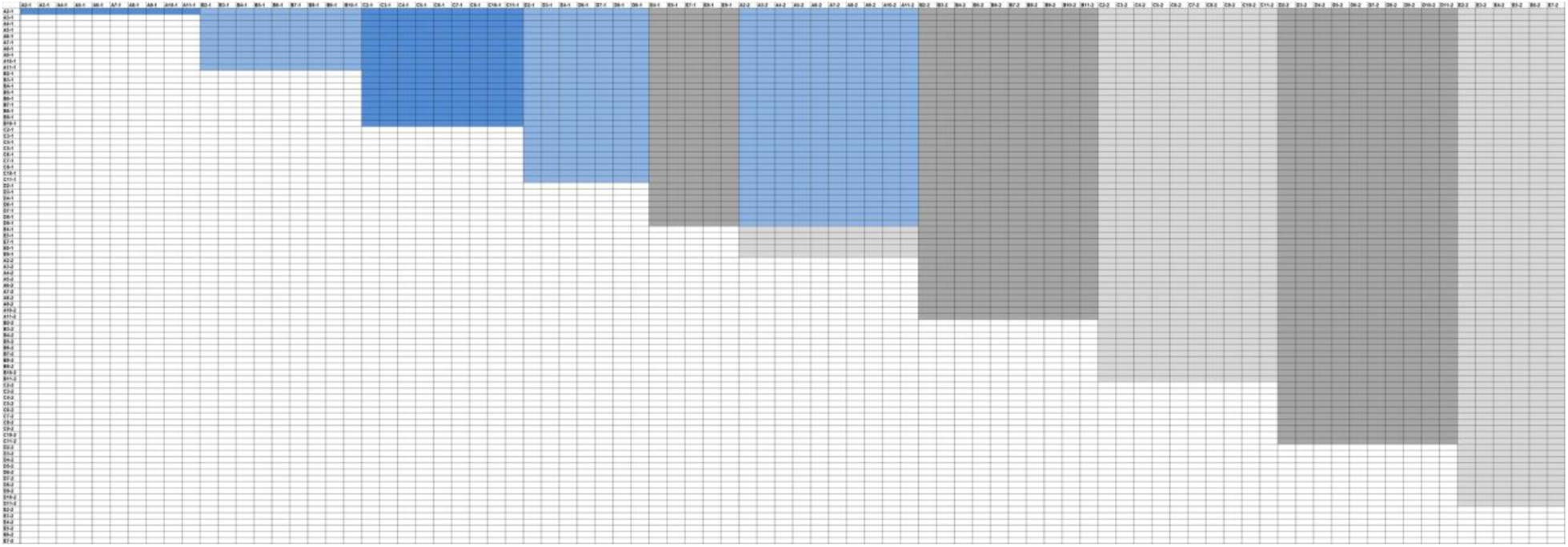
Grid representation of the Cayman Chemical Epigenetic Screening Library drugs, in which each drug (86 out of 94 total library drugs) is represented along both the x- and y-axes. See Table 1 for the list of drugs. Table 1 drugs that are removed from this grid representation are CAY10669, GSK-J1, GSK-J4, Tenovin-1, (-)-JQ1, MI-nc, GSK-J2, and GSK-J5. Shading (dark blue/light blue, dark grey/light grey) represents rows of drugs in the Library plates. Each horizontal, alternately-shaded row represents a candidate drug pool. Blue shading represents the 93 pools tested. Grey shading represents the remaining candidate pools (393 total on grid).

